# Flies exploit predictable perspectives and backgrounds to enhance iridescent signal salience and mating success

**DOI:** 10.1101/733758

**Authors:** Thomas E. White, Nina Vogel-Ghibely, Nathan J. Butterworth

## Abstract

Communication requires both the encoding of information and its effective transmission, but little is known about display traits that primarily serve to enhance efficacy. Here we examined the visual courtships of *Lispe cana*, a cursorial fly that lives and mates in heterogeneous foreshores, and tested the prediction that males should seek to enhance signal salience and consequent fitness through the flexible choice of display locations. We show that courting males access the field of view of females by straddling them and holding their wings closed, before moving ahead to present their structurally coloured faces in ritualised dances. Males preferentially present these UV-white signals against darker backgrounds, and the magnitude of contrast predicts female attention, which in turn predict mating success. Our results demonstrate a striking interplay between the physical and attentional manipulation of receivers and reveal novel routes to the enhancement of signal efficacy in noisy environments.

## Introduction

Communication requires the effective transmission and reception of information in complex natural environments. Selection has favoured diverse solutions to this basic challenge, which are showcased among the visual ornaments and displays of animals (Dalrymple et al. 2018; Maia et al. 2013; Girard et al. 2015; White 2018). The functions of traits involved in sexual communication are twofold. Namely; to encode information relevant to mate choice and assessment (i.e. signal content) and ensure its effective transmission and reception (i.e. signal efficacy). Extensive work on content has shown how colour signals and (to a lesser extent) behaviours can encode information on benefits to potential mates, and are the targets of pre-copulatory choice (Brooks and Endler 2001; Kemp 2008a; Kemp 2008b; Barry et al. 2015). This is well illustrated in the literature on pigmentary colour expression and individual condition. The production of carotenoid-based signals, for example, can be physiologically tied to past or present body condition, reproductive and parental quality, and/or immune function, and so offer an honest guide to aspects of mate quality (Weaver et al. 2018). Less is known, however, about the features of displays that act primarily in the service of efficacy.

Most signalling traits are multifunctional (Candolin 2003; Bro-Jørgensen 2010). In addition to encoding information, colour patterns and displays may capture attention (Ord and Stamps 2008), amplify the conspicuousness of other traits (Smith et al. 2009), or modify the content of co-expressed signals (Endler et al. 2014). Traits that serve such purposes are not the target of choice but are nonetheless key determinants of attractiveness. This is particularly true in heterogeneous environments which will modify, and set limits on, the salience of signals, and the outcomes of selection for efficacy under such conditions are broadly predictable. In evolutionary terms, signal forms that best cut through noise to stimulate receivers will be favoured (Dusenbery 1992). Structural colours, which arise from the selective reflection of light by nano-scale structures, offer one such solution since they are capable of generating uniquely broad, rich, and dynamic colour palettes (Greenewalt et al. 1960; Vigneron and Simonis 2010; White et al. 2012; Maia et al. 2013), and are evolutionarily labile (Maia et al. 2013; Wasik et al. 2014). Of particular interest to this study is their potential for extreme chromaticity and brightness, which are valuable for maximising basic features of efficacy such as signal conspicuousness (Schultz et al. 2008) and detectability (Schultz and Fincke 2009) in the wild. In ecological terms, environmental complexity can drive flexible behavioural solutions that improve the signal-to-noise ratio. These include the precise behavioural delivery of signals (White et al. 2015; Simpson and McGraw 2018), varying the timing and duration of displays (Poesel et al. 2006), and selecting optimal locations for courtship (Endler and Thery 1996).

Flies possess a suite of colour-producing mechanisms and display behaviours and so offer excellent, albeit underutilised, opportunities for exploring questions of signal evolution (Marshall 2012). The genus *Lispe* is a cursorial group of muscids that spend much of their life on or near the ground in littoral habitats (Pont 2019). This includes courtship, during which males pursue and physically straddle walking females before presenting their iridescent faces and wings in ritualised ‘dances’ (Frantsevich and Gorb 2006; Pont 2019). Though excellent work continues to document the structure of these and related displays (Spofford and Kurczewski 1985; Frantsevich and Gorb 2006; Jones et al. 2017; Butterworth et al. 2019), it is not known whether or how such colours and behaviours serve to effectively transmit information to mates. Given that the visual structure of seaweed-dominated foreshores will vary over short temporal and spatial scales (e.g. with tides), theory predicts functional links between signal structure and display behaviours to enhance signal salience (Lythgoe 1979; Dusenbery 1992). Specifically, selection should favour male colour traits that are reliably conspicuous to conspecifics in their surrounds, and/or which are delivered via flexible behaviours for exploiting locally optimal conditions.

Here we examine the courtship displays of the fly *Lispe cana* with a view to testing how signal structure and display behaviour mediate communication efficacy and mating success. *Lispe cana* is a species endemic to Australia that inhabits supralittoral (foreshore) habitats spanning the east coast. Males and females have striking structurally-coloured ‘white’ and ‘yellow’ faces (Fig. 1), respectively, that are weakly iridescent across lateral viewing angles and more strongly so in the dorsevental plane (unpublished data). They are active predators of other ground-dwelling invertebrates and they live, hunt, and mate on shorelines (ca. 0-5 m from the waterline) which are often populated by patchy distributions of seaweed and detritus (Fig. 1). The casual observation of courtship in this species shows that smaller (5.5 - 7 mm body length) males approach and straddle the larger (6.5 - 8 mm) females from behind, and hold the female’s wings closed using their forelegs (Fig. 2 a-b; supplementary video S1). Males maintain this position as females continue to move about the environment. After a time, males rapidly move in front of females and present their iridescent faces and wings in ritualised displays that consist of erratic movements around the female’s head at very close distance (Fig. 2c). These displays then typically terminate when females lose interest and disperse, or when males re-mount and mate with receptive females (Fig. 2d). This presents ecologically tractable opportunities to examine the predicted links between signal structure and signalling behavior in the service of efficacy, since the initial straddling of mates implies the potential for males to optimise the timing and location of displays given their largely shared field of view. In a field-based assay, we thus quantified the visual structure of signals and signalling environments and tested the prediction that males should seek to maximise the conspicuousness of their facial signals within their dynamic visual habitats to enhance mating success.

**Figure 1:**
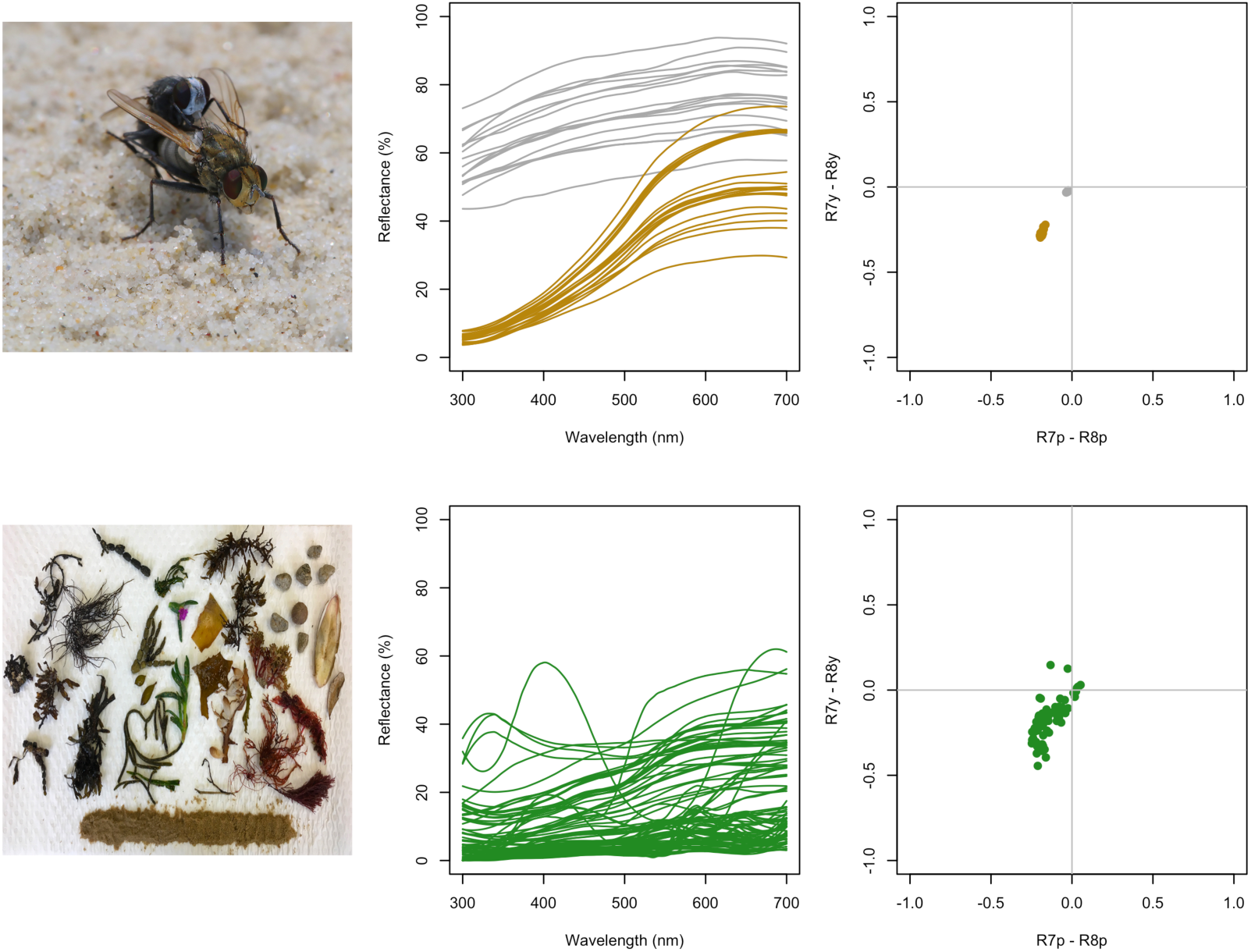
Male and female *Lispe cana* and material sampled from their foreshore habitats (left; photo credits NB & TEW), the reflectance spectra of male (grey) and female (yellow) faces and background material (centre), and the spectra as represented in a muscid colourspace (right).

**Figure 2:**
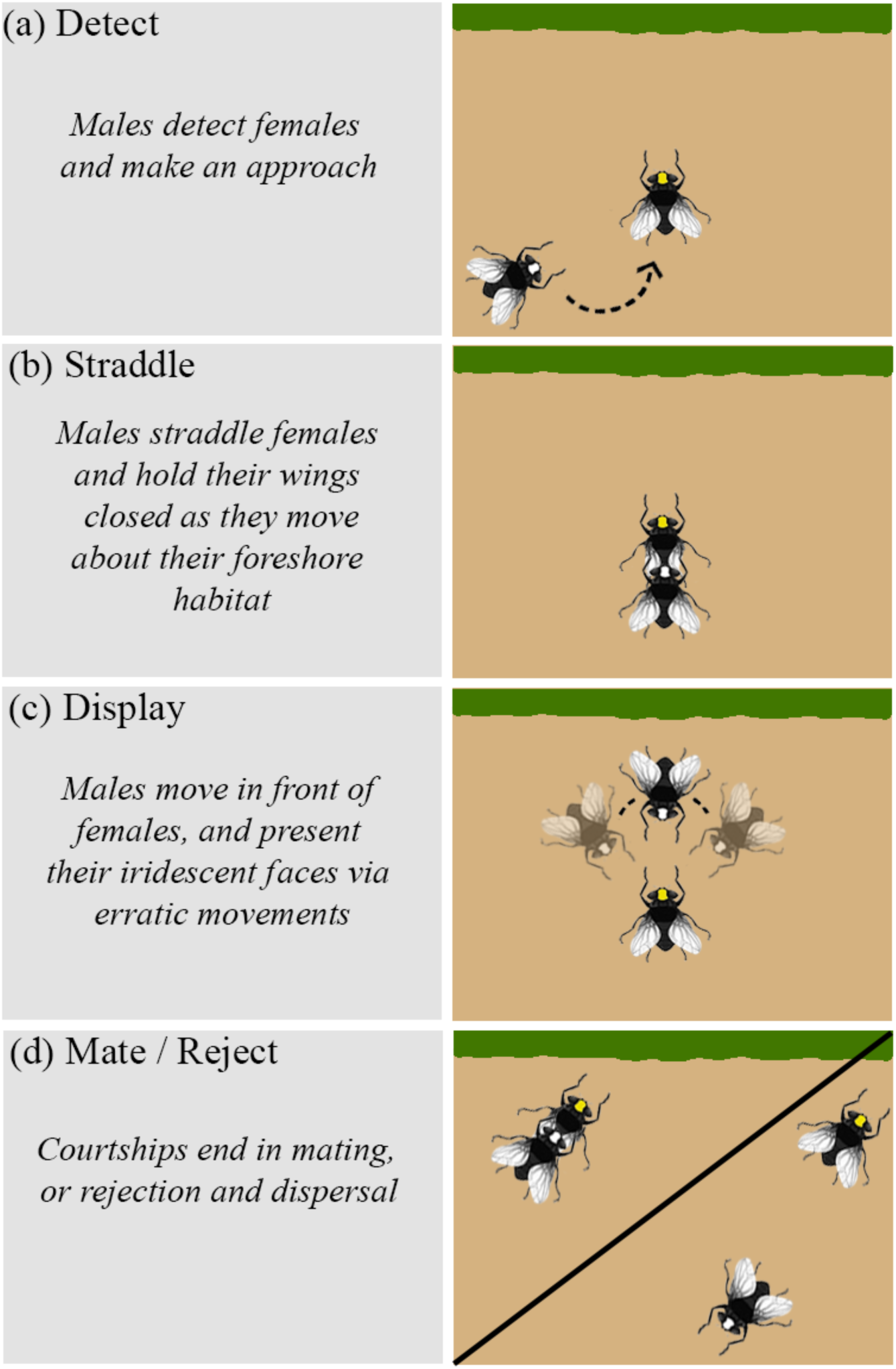
Courtship in the cursorial, shore-dwelling muscid fly *Lispe cana*. Male *L. cana* detect females within their seaweed-dominated habitats and rapidly approach (a), before ‘straddling’ females from behind and holding their wings closed (b). During this time males are aligned with females (2.1 ± 1.4°offset, 29.4 ± 7.5°elevation), and are thus sharing a field of view. Males then preferentially display against backgrounds that enhance the luminance contrast of their UV-white faces, which leads to improved salience, female attention and subsequent mating success.

## Methods

### Sampling courtships, flies, and visual backgrounds

We recorded 42 independent courtships of *L. cana* using a GoPro Hero 6 at either 30 or 60 fps in the field. From these we extracted the duration of the straddling and display phase of each courtship, which are readily identified by eye (Fig. 2b-c). We also estimated the male-female alignment and elevation of straddling males in three haphazardly selected frames from each courtship sequence, which we averaged. Alignment was taken as the angle between male and female midlines (e.g. with 0 indicating complete alignment) as measured from points on the centre of their heads and abdomens from top-down video. Similarly, male elevation was taken as the angle between male and female midlines from a lateral view. We were unable to extract complete data for nine courtship sequences due to a lack of suitable viewing geometries.

All courtships were recorded within a 6 m^2^ region of the supralittoral zone of Toowoon Bay, NSW, Australia, on clear days between 1100 and 1300. We used a fixed region for our observations so that we could subsequently sample the entirety of the visual environment for analysis (detailed below), and restricted the observation period to hold the sun’s azimuth approximately constant, and so minimise any effects of light-source directionality on variation in signal appearance and display behaviour. Flies typically dispersed outside the observation area at the conclusion of a display, minimising the risk of repeated recordings, and the few that did remain were not intentionally observed again. Following each display, we noted whether the interaction resulted in copulation, and collected a sample of the nearest piece of background material within 150 mm directly behind the courting male. If no such material was present, we took sand to be the relevant visual background. At the conclusion of the experiment we also gathered all material within the 6 m^2^ observation area as a representative sample of the all possible visual backgrounds available to males. From these we collected reflectance measurements at three evenly spaced points along the length of each piece of material using the general method described below and averaged them. We also photographed the focal area at the conclusion of the experiment and estimated the proportional area of open sand using Adobe Photoshop CC (v20.0.6), before collecting haphazardly-selected samples for spectral measurement. By systematically collecting and measuring all material within the area of observation, and including spectral samples of sand proportional to its availability, we also approximately accounted for the relative abundance of potential viewing backgrounds in our analyses of background selection and bias (detailed below).

We also separately collected 17 male and 20 female flies and recorded the reflectance of their facial colouration to calculate a population-average estimate of facial colouration for use in our analyses of signal/background contrast. This was necessary because the rapid dispersal of flies at the conclusion of courtship meant we were unable to sample the observed courting pairs described above. We used an OceanOptics Jaz UV-VIS spectrometer with a pulsed xenon light source, set to an integration time of 50 ms with a boxcar width of five. We used a 400 um bifurcated probe oriented normal to the plane of each fly’s face to capture the entirety of its ca. 5 mm^2^ area, and recorded and averaged two measurements per individual. When measuring background material, we instead used a measurement geometry of 0° illumination and 45° collection to minimise specular reflections from damp surfaces. A spectralon 99% diffuse reflector (Labsphere, New Hampshire) and black velvet served as our light and dark standards, respectively, and we recalibrated between each measurement. Finally, we binned the resulting spectra at 1 nm intervals and applied some minor locally weighted scatterplot (LOESS) smoothing (*α* = 0.15) prior to analysis.

### Visual modelling

We used an early-stage (i.e. retinal-level) model of muscid colour vision to estimate the subjective colour and luminance contrasts of male *Lispe* faces against presentation backgrounds. We drew on the receptor sensitivities of *Musca domestica* as the closest available relative of *Lispe*, and assumed the involvement of R7p, R8p, R7y, and R8y photoreceptors in chromatic processing, and R1-6 receptors in achromatic processing (Hardie 1986; Troje 1993). For chromatic contrasts, we drew on the model of Troje (Troje 1993) and estimated receptor quantum catches as the integrated product of receptor sensitivity, stimulus reflectance, and a standard-daylight illuminant. We then calculated the difference in relative stimulation between R7y-R8y and R7p-R8p receptors as putative opponent channels that define the location of a given stimulus in dipteran colourspace (Fig. 1, right side), and chromatic contrasts were taken as the Euclidean distance between stimuli in this space. We estimated luminance contrast as the Weber contrast of fly faces against their backgrounds, with quantum catches calculated as above albeit using the R1-6 receptor absorbance. All spectral processing, analysis, and visual modelling was carried out using the package package ‘pavo’ v2.2 in R v3.5.2 (Maia and White 2018; R Core Team 2018; Maia et al. 2019).

### Statistical analyses

We first tested whether male flies display against visual backgrounds non-randomly with respect to their signal contrast. We used a bootstrap test which, for a given run, entailed drawing 42 backgrounds with replacement from the total pool of background material collected from the observation area, before calculating the mean of the chromatic and achromatic contrasts of the average male face against each item. We repeated these 5000 times, thereby generating a null distribution of contrasts that represent the probability of observing a given mean chromatic or achromatic contrast value under the assumption that males display against backgrounds at random. We then calculated the probability of attaining our observed chromatic and achromatic contrast values given this null expectation and used Cohen’s d (the difference between sample and null means, divided by the pooled standard deviation) as an estimate of the magnitude of any difference between observed and null-distributed contrast values.

We then examined the effects of signal contrast and two proxy measures of female attention on mating success using generalised linear models in a restricted maximum likelihood information-theoretic framework (Anderson and Burnham 2004). We took the duration of male displays as a between-courtship measure of attention, because when males release females and move to present their facial signals (Fig. 2b-c) the duration of the subsequent display is largely under female control. Females are free to disperse at any point during the male display and so terminate it, which is indeed the most common outcome of a courtship interaction (Fig. 2d). Our second, within-courtship, measure of female attention drew on the fact that females repeatedly re-orient themselves in brief, saccadic movements during male displays (supplementary video). We therefore calculated the proportion of these reorientations that were directed toward displaying males, as defined by a reduction in the angle between the female midline and male head. A value of one indicates that all female movements were toward males during their displays, while a value of zero means that females consistently oriented away from displaying males. Of course, these are only approximate measures, but the above rationale, and the correlation of both measures (Pearson’s r = 0.33, t_40_ = 2.22, p = 0.03), suggest that they collectively capture relevant aspects of female attention.

We specified a global model with mating success as a binomial response (with logit link function), and included male display duration, the proportion of female reorientations toward displaying males, and chromatic and achromatic signal contrast as main effects. We examined all models containing linear combinations of these predictors along with an intercept-only null model and ranked them according to Akaike’s Information Criterion (AICc, adjusted for small sample sizes). We calculated the R^2^ for the most parsimonious model(s), as approximated by a *Δ*AICc of < 2 (Anderson and Burnham 2004).

We also examined the effect of signal contrasts and straddling duration on both the duration of male displays, and the proportion of female orientations toward displaying males (i.e. female attention). We modelled display duration as a Gaussian response (with identity link function) and female orientations as a binomial response (with logit link) and included chromatic and achromatic signal contrasts and the duration of the straddling phase as main effects in both models. We visually confirmed the assumptions of normality among residuals and homogeneous variance structures for all models and used the package ‘MuMIn’ v1.43.6 (Bartoń 2013) for all model selection in R (R Core Team 2018).

## Results

During courtship male *L. cana* pursue lone females before straddling them and holding their wings closed with their forelegs (Fig. 2). While in this position, males are slightly elevated (29.4 ± 7.5°) and closely aligned with the body axis of females (2.1 ± 1.4°), and are afforded direct access to the female’s field of view (Fig. 2b). This straddling phase lasted 1.58 - 16.53 seconds, after which females were released by males who then repeatedly ran in close semi-circles in front of females as part of a ritualised display (supplementary video; also see Frantsevich and Gorb 2006; Butterworth et al. 2019 for comparable examples). During these display phases which lasted 0.76 - 11.60 seconds, male *L. cana* preferentially presented themselves against backgrounds that generated strong achromatic (p < 0.001, d = 3.363) and chromatic (bootstrap p = 0.047, d = 1.744) contrast with their facial colouration (Fig. S1). That is, their faces appear much brighter and slightly more colourful to conspecifics than would be expected if they were displaying at random within their visual environments.

The most parsimonious model of mating success (12/42 courtships) indicated a strong positive contribution of both of our included measures of female attention; the total duration of male displays, and the proportion of female orientations toward displaying males (Table 1; Table 2). It clearly outperformed the null ((*w*_1_ + *w*_2_)/*w*_null_ > 700), and was approximately three times as informative as the second and third best models that included the additional effects of chromatic contrast (*Δ*AICc = 2.15, *w*_1_/*w*_2_ = 2.93) and achromatic contrast (*Δ*AICc = 2.44, *w*_1_/*w*_3_ = 3.38), respectively. This suggests that female attention (as measured via male display duration and female orientations) directly mediates mating success, which is further supported by the predictive relationship between achromatic, but not chromatic, signal contrast and both display duration and female orientations (Table 3; Fig. 4). The effect of signal/background luminance contrast on mating success is thus almost entirely driven by its influence on female attention. That is, males which presented more contrasting signals through the selective use of backgrounds were able to display longer and better held females’ focus, which improved mating success. Finally, we found no effect of straddle duration on display duration or female orientations (Table 4), indicating that the duration of the two courtship phases is unrelated and, more broadly, that female receptivity during straddling and display phases is unrelated.

**Table 1:**
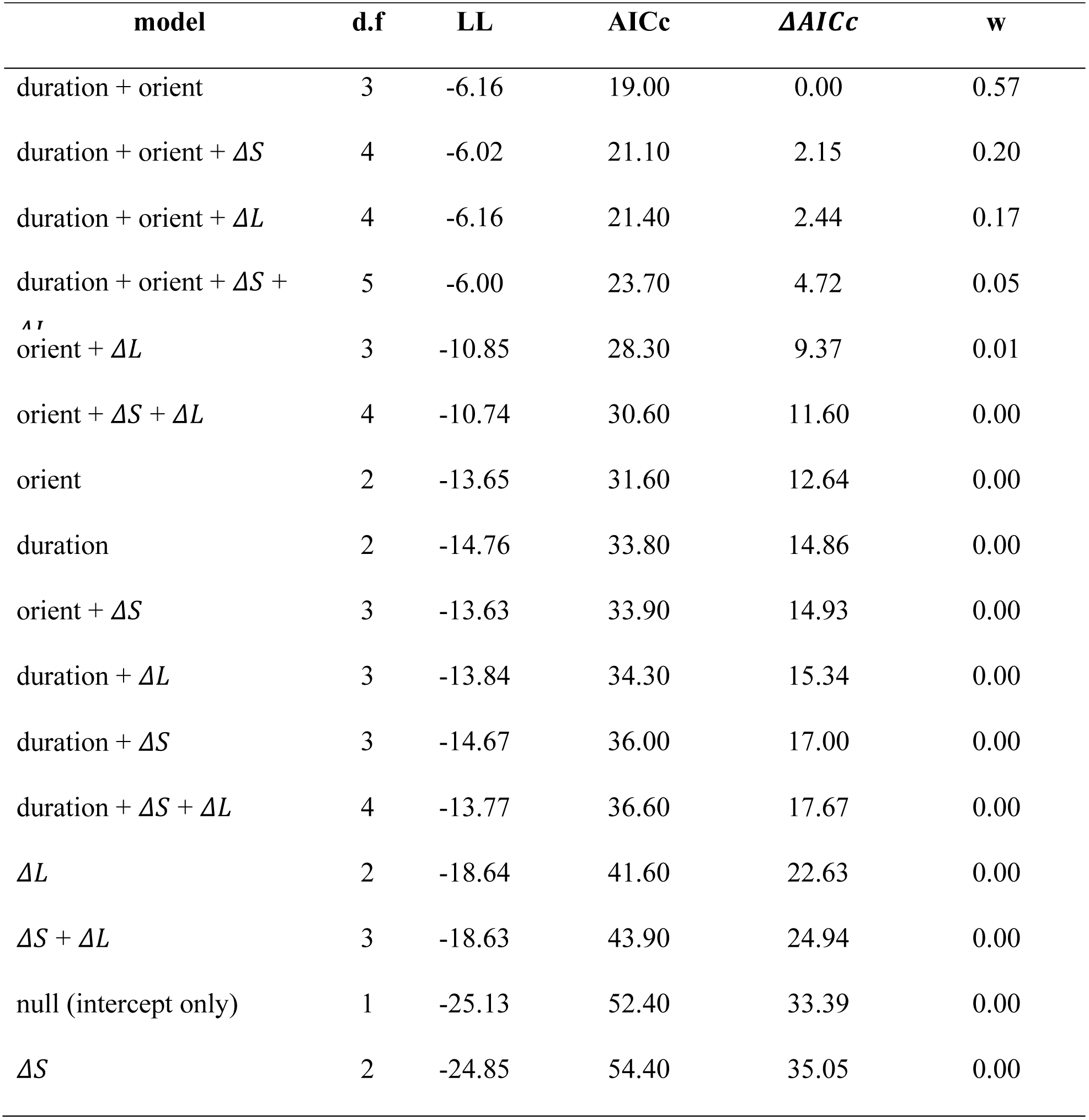
Full model selection table, detailing the relative strength of candidate binomial GLM’s for the relationship between mating success and one or more linear combinations of courtship display duration (dur), the proportion of female orientations toward males during displays (orient), and the chromatic (*Δ*S) and achromatic (*Δ*L) contrast of male facial colouration against their visual backgrounds.

**Table 2:** Full results of the leading model of mating success that includes the effects of male display duration and the proportion of female orientations toward males during their displays, drawn from a global binomial GLM that included display duration, female orientations, chromatic contrast, and achromatic contrast as predictors (Table 1).

| parameter | estimate | std. err. | z | p | R<sup>2</sup> |
| --- | --- | --- | --- | --- | --- |
| intercept | -12.93 | 4.72 | -2.74 | < 0.001 | 0.68 |
| display<br>duration | 1.14 | 0.54 | 2.10 | 0.035 |  |
| orientations | 9.61 | 3.67 | 2.62 | < 0.001 |  |

**Table 3:**
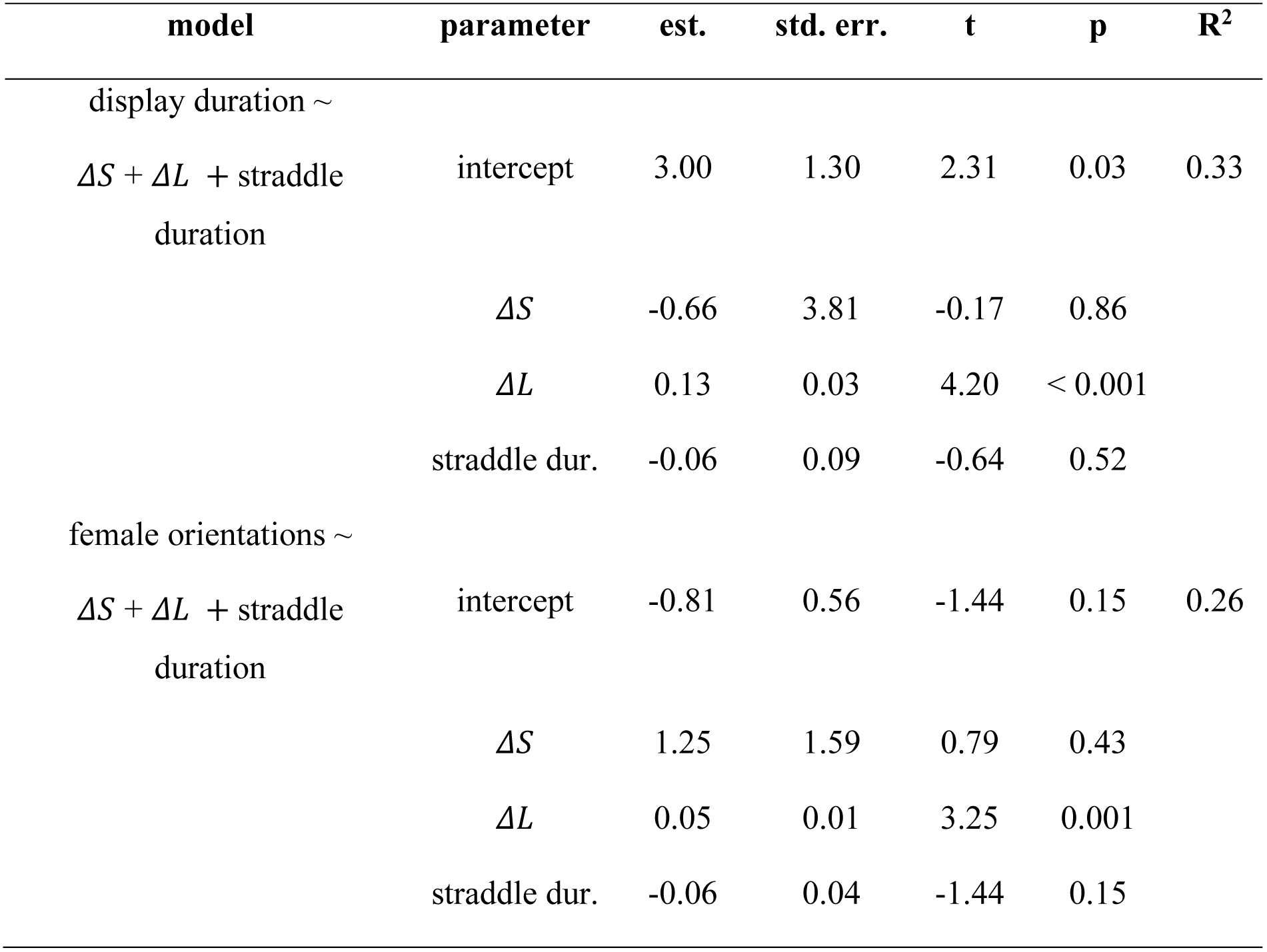
Results of separate GLM’s estimating the effect of chromatic (dS) and achromatic (dL) face/background contrasts, and the duration of the ‘straddling’ courtship phase on two proxy measures of female attention: the duration of male *L. cana*’s courtship displays, and the proportion of female orientations toward males during their display.

| model | parameter | est. | std. err. | t | p | R<sup>2</sup> |
| --- | --- | --- | --- | --- | --- | --- |
| display duration ~ |  |  |  |  |  |  |
| $\Delta S + \Delta L + \text{straddle}$<br>duration | intercept | 3.00 | 1.30 | 2.31 | 0.03 | 0.33 |
| | $\Delta S$ | -0.66 | 3.81 | -0.17 | 0.86 | |
| | $\Delta L$ | 0.13 | 0.03 | 4.20 | < 0.001 | |
|  | straddle dur. | -0.06 | 0.09 | -0.64 | 0.52 |  |
| female orientations ~ |  |  |  |  |  |  |
| $\Delta S + \Delta L + \text{straddle}$<br>duration | intercept | -0.81 | 0.56 | -1.44 | 0.15 | 0.26 |
| | $\Delta S$ | 1.25 | 1.59 | 0.79 | 0.43 | |
| | $\Delta L$ | 0.05 | 0.01 | 3.25 | 0.001 | |
|  | straddle dur. | -0.06 | 0.04 | -1.44 | 0.15 |  |

**Figure 3:**
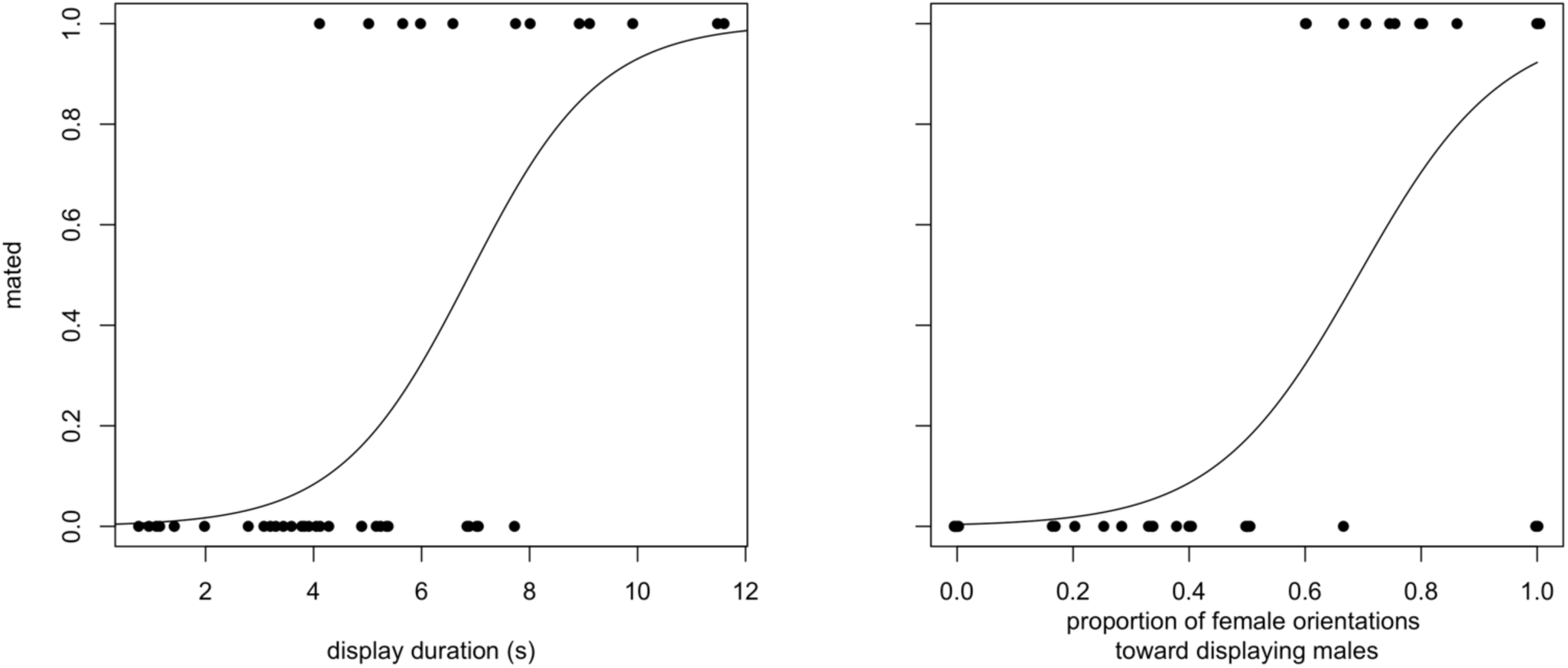
The relationship between two measures of female attention — display duration and female orientations toward male displays — and mating success in *Lispe cana*. Circles represent the presence or absence of mounting following courtship, with the line indicating model fit as estimated by a binomial GLM (Table 2).

**Figure 4:**
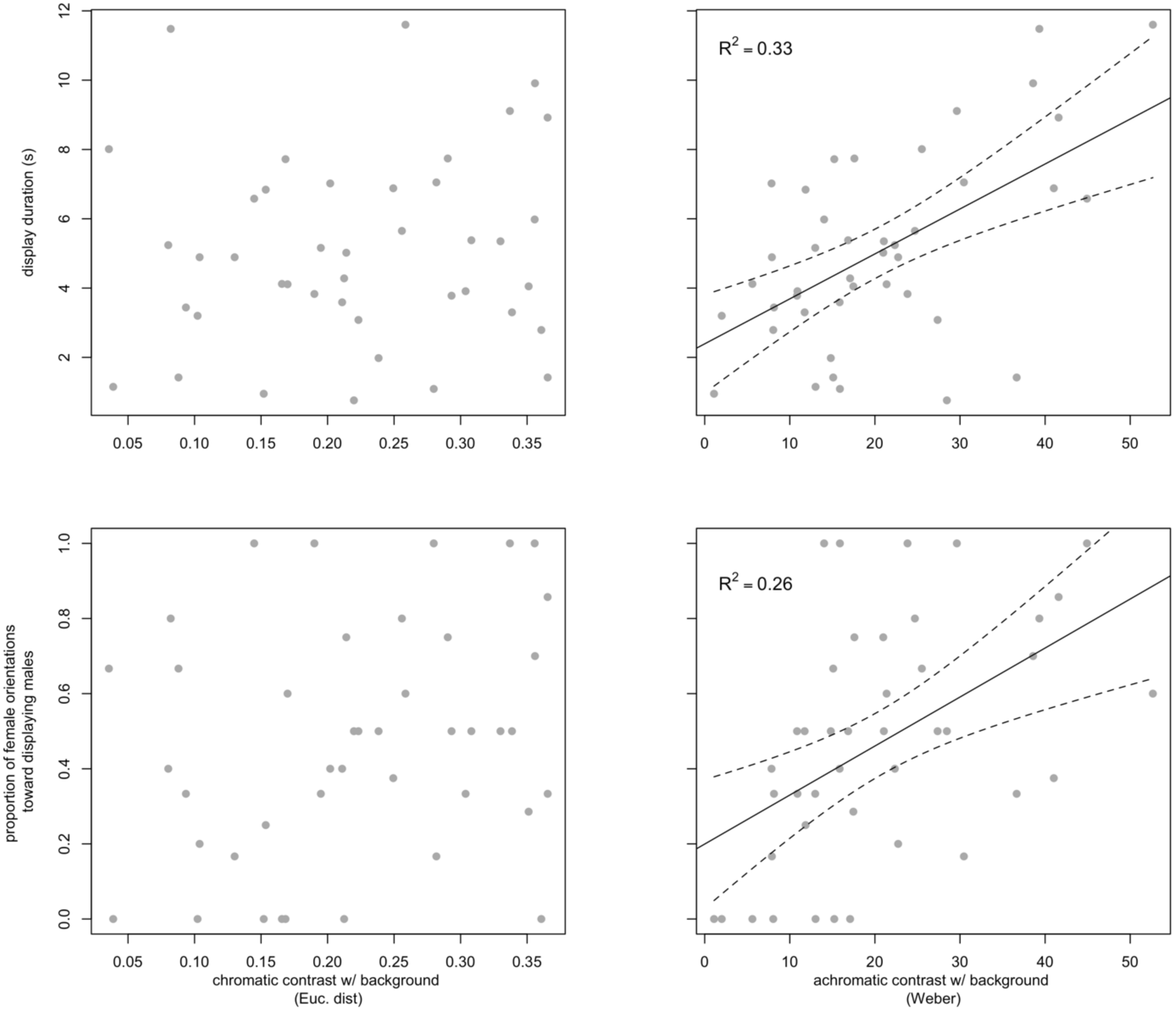
The relationship between the chromatic and achromatic facial/background contrast of male *Lispe cana*, male display duration, and female orientations toward displaying males. Solid and dashed lines indicate the best fit ± 95% CI as estimated by a GLM (table 3).

## Discussion

Selection has generated myriad solutions to the challenge of communicating amidst noise, with animals varying the location (Endler and Thery 1996), timing (Poesel et al. 2006), and behavioural delivery (White 2017; Simpson and McGraw 2018) of signals to ensure their effective reception. Here we reveal novel links between a structural colour signal and its behavioural presentation, which in part relies on the exploitation of the receiver’s predictable field of view. Male *Lispe cana* straddle and align themselves with females during courtship, before releasing and displaying their conspicuously coloured faces in a ritualised dance (Fig. 2). Males preferentially display against darker backgrounds which enhances visual contrast (Fig. S1), leading to longer display times overall and increased female attention toward male signals (Fig. 4) and, in turn, enhanced mating success (Fig. 3). Our results suggest that males are flexibly exploiting females’ perspectives and visual environments to enhance the attractiveness of their complex displays and, hence, mating success.

The courtships of *L. cana* reveal a striking interplay between the physical and attentional manipulation of receivers. When straddling, males physically impede females from flying, and the independence of straddling and display durations suggests that it proceeds until either (the consistently larger) females forcefully disengage males, or males release to display (Table 1). Whether males have any control over the mobility of walking females, or present an encumbrance at all, is unclear, though the near-constant movement of females (pers. observation) means that diverse display locations may be passively sampled within a short time. Once males do release then females are physically free to leave, which necessitates the capturing of female attention during signalling to ensure further receptivity and mating. The predictive relationships between facial contrast and female attention (Fig. 3), and female attention and mating success (Fig. 4), suggest a central role for male signals and presentation behaviours in maintaining the focus and receptivity of females amidst cluttered visual environments. The structurally generated UV-white colour of male faces will, by definition, maximise luminance contrast among the desaturated hues of surrounding seaweed (Fig. 1; though less so among sand), and their flexible display behaviours, combined with predictable knowledge of viewer perspectives, allow for the exploitation of locally optimal conditions. Thus, males presenting greater luminance contrast are afforded longer displays by viewing females as well as greater female attention during their displays, with tangible benefits for mating success.

Several aspects of male displays are likely to further improve salience. For one, the rapid movement of males from outside the female field of view to within it between straddling and display phases (Fig. 2b-c) will generate strong temporal contrast. This may be amplified by the adaptation of female receptors to visual backgrounds immediately prior to seeing the male display, since the fastest component of receptor adaptation takes less than a second (Smirnakis et al. 1997; Shevell 2001; Baccus and Meister 2002). The continual, erratic movement of males during displays will also sweep out a larger area of the female’s retina (supplementary video; unpublished data), further enhancing its salience. Analogous display features are described in male great bowerbirds *Ptilonorhynchus nuchalis*, which offer an illustrative comparison. Males construct bowers and exploit the fixed perspective of females to induce a visual illusion, the quality of which is predictive of male mating success (Kelley and Endler 2012). In addition to this focal display males use foraged materials to colour the walls of their avenues and also actively flash coloured objects across the female’s field of view. Though not associated with mating success directly, these latter features serve to increase communication efficacy through the capturing of receiver attention and enhancement of other signals (Kelley and Endler 2012; Endler et al. 2014).

The strong effects of display duration and female orientations on mating success suggests that females may be assessing content-rich features of the male display, since the informational load of a task increases decision times (Chernev et al. 2015; Hemingway et al. 2019). Though not directly explored here, the structurally coloured faces of male *L. cana* are theoretically well-suited conduits of information on mate quality. This colouration is primarily a consequence of light scattering by flattened bristles (Frantsevich and Gorb 2006), which demand close developmental control during metamorphosis for optimal expression (Ghiradella and Butler 2009). Variation in the resulting signal may thus inform potential mates about foraging ability, developmental stability, and/or aspects of genetic ‘quality’ (Kemp 2008b; Barry et al. 2015). The primacy of luminance over chromatic contrast in our tests (Table 1; Table 2) suggests a role for signal brightness and variation therein as the more salient channel. It is also consistent with the relatively under-developed colour sense of flies, which instead tend to draw on luminance to guide the identification and categorisation of stimuli (Troje 1993; Lunau 2014). Though luminance is a less reliable cue than hue in natural environments, it is a known channel of sexual communication in several insect species (Kemp 2008a; Kemp 2008b; Barry et al. 2015). Well-described examples include the mantid *Pseudomantis albofimbriata* in which the brightness of females abdomens is tied to their current condition and is the focus of male choice (Barry et al. 2015), and the butterfly *Eurema hecabe*, wherein the intensity of male UV wing colouration encodes information on larval resource acquisition (Kemp 2008b; Kemp 2008a).

Despite the plausibility of *L. cana*’s faces as informative signals, we found no direct relationship between visual contrast and mating success as might be predicted for traits under selection for such a purpose. There are several reasons for this. For one, the role of any given trait may be to hold attention, as our results show is at least partly the case (Fig. 4), with others acting as more direct indicators of mate quality. The iridescent wing interference patterns of male *L. cana* that are actively presented during displays offer one such candidate trait, and emerging evidence suggests these patterns are both widespread among flies and may be subject to sexual selection (Katayama et al. 2014; Hawkes et al. 2019). A closely related possibility is that male facial colouration is an amplifier (Hasson 1991; Byers et al. 2010), which serves to make the direct targets of choice easier to discern or assess. Alternately, if male facial colouration is instead a signal of species or sex recognition, as suggested by the striking sexual dimorphism (Fig. 1) and variation in facial colouration among sympatric species (Pont 2019), then female preferences may be expressed as a simple threshold function. That is, the trait may have to be above a certain value to be effective, but variation beyond that will be irrelevant. Finally, and more specific to this study, is the fact that we averaged facial reflectance across a sample of flies that were not observed in the focal courtships themselves, which prevents us from assessing the direct contribution of individual-level signal variation (in addition to behavioural variation) to mating success.

Environments frequently modify, and set limits on, the salience of sexual displays. Here we describe an innovative solution to this problem that relies on the physical and attentional manipulation of viewers. By adopting the perspective of receivers during courtship, male flies are able to select locally optimal display locations to enhance the salience of their sexual signals and consequent mating success. More broadly, our results suggest that conspicuous signals and flexible display behaviours can arise in response to environmental heterogeneity. Display site properties will vary between habitats, however, which may mediate local adaptation and sexual isolation. This will be most pronounced between habitats that vary acutely in relevant aspects of structural complexity, such as the reliably seaweed-laden foreshore of our focal population, and the entirely barren beaches of nearby populations. This presents intriguing opportunities for illuminating the role of sexual communication in diversification, for which tractable groups such as *Lispe* hold excellent promise.

## Acknowledgements

TEW thanks Elizabeth Mulvenna and Cormac White for their support. TEW also thanks Hannah Rowland for the salient observation of parallels between *Lispe* and bowerbirds, which sparked closer consideration of a fly’s view of courtship. We appreciate the thoughtful comments of Professor Edmund Brodie and two anonymous reviewers, whose input has materially improved the clarity and robustness of this work.

## Data availability

All data and code will be made persistently available upon acceptance.

